# Antimicrobial, Antioxidant and Anti-Inflammatory Activities of Proteins of *Phaseoulus lunatus* (Fabaceae) Baby Lima Beans Produced in Ecuador

**DOI:** 10.1101/401323

**Authors:** J. Tamayo, T. Poveda, M. Paredes, G. Vásquez, W. Calero-Cáceres

## Abstract

*Phaseoulus lunatus* L., a variety of baby lima bean, which is produced in the coastal region of Ecuador, is a profitable crop of that country. Various cultivars of this common bean are considered a sources for nutraceutical compounds, such as bioactive peptides. To assess the potential biologic activities of protein isolates and hydrolysates of *P. lunatus* baby lima beans, this study evaluates the proteins antimicrobial, antioxidant and anti-inflammatory activities. Antioxidant activity was measured by the TBARS method. In-vitro anti-inflammatory activity was measured by the inhibition of denatured protein as well as a diffusion method, according with CLSI guidelines by antimicrobial activity. Both fractions (isolate and hydrolysates) showed anti-inflammatory and antioxidant activity. However, protein hydrolysates (pH 5) had a better performance than protein isolates. The same effect was observed in antimicrobial activity, when protein hydrolysates had a broad-spectrum antimicrobial activity against Gram-negative and Gram-positive bacteria. These preliminary studies suggest that *P. lunatus* baby lima beans could have a considerable biological activity for nutraceutical applications.

## 1 INTRODUCTION

The genus *Phaseolus* (beans), included in the family *Leguminosae*, represent one of the most ancient, common legumes consumed by humans around the world. In developing countries, this legume specie is one of the principal sources of dietary proteins (BROUGHTON *et al.* 2003). Today, in terms of morphological and genetic diversity, legumes are a complex genus with four genetic groups (BITOCCHI AND NANNI 2012). Previous research has shown that the northwestern region of South America (Colombia, Ecuador and northern Peru) represents the convergence of the two principal genetic *P. vulgaris* varieties (Central American and Andean) (DEBOUCK *et al.* 1993). This convergence shows a clear difference of morphology and molecular characteristics. Considering the current correlation between population growth and bean consumption (BELLUCCI *et al.* 2014), a detailed analysis of chemical and biological characteristics would help us better understand the properties and differences between common bean varieties.

An increasing amount of literature demonstrates the biological properties of protein isolates of legumes. These properties represent a considerable nutritional role and a source of essential amino acids (BETANCUR-ANCONA *et al.*, 2009; CARBONARO, MASELLI, & NUCARA, 2015). Varieties and cultivars of common beans has been considered as sources of nutraceutical compounds, such as bioactive peptides, polyphenols, and polysaccharides (NGOH *and* GAN 2016; CAMPOS-VEGA *et al.* 2010; BETANCUR-ANCONA *et al.* 2009; BITOCCHI *and* NANNI 2012).

The variety of baby lima bean that is produced in the coastal region of Ecuador is a profitable crop for that country in terms of economic benefits (EL PRODUCTOR 2015). At the moment, no published information about physicochemical and biological characteristics of this Ecuadoran variety of bean. Considering the differences between *Phaseolus* germplasm according their precedence (BERNAL *and* GRAHAM 2001), the differences are expected in chemical composition and biological activities. This study evaluates the antimicrobial, antioxidant and anti-inflammatory activities of protein extracts and protein hydrolisates of *P. lunatus* var. baby lima beans, in order to identify the beans potential source of bioactive compounds for nutraceutical applications.

## 2 MATERIALS AND METHODS

### 2.1 Samples

*P. lunatus* var. baby lima beans were purchased in a local market in Machala, Ecuador. Seeds were dehydrated in a Laboratory Incubator (ESCO, Singapore) at 50 °C for 48 hours. Subsequently, dehydrated seeds were grinded into a fine flour powder with a mill MG 1511.

### 2.2 Protein isolates from baby lima beans

Baby lima bean protein concentrate was obtained according to the methodology of BARRIO & AÑÓN, (2010) with minor modifications. Homogenized flour was diluted in a 1:10 solution using distilled water. The pH level was set at 8 by using sodium hydroxide 2 M for 1 hour, shaken at 800-900 rpm. Those samples were centrifuged at 4.400 rpm for 30 minutes. The supernatant was recovered and treated at different pHs (3, 4, 5 and 6) using, if necessary, hydrochloric acid 2 N or sodium hydroxide 2M. The mixture was vigorously homogenized and stored at 4 °C for 24 hours. The supernatant was discarded and the precipitate**s** were freezed at −80 °C and lyophilized using a freeze dryer Bench Top Pro BTP-3ES0VW (SP Scientific, Stone Ridge, NY, USA), at −50 °C and 0.2 Pa. The extraction performance was calculated with this formula: % performance = (Pf / Po) * 100. Where Po: Initial weight of flour, Pf: Final weight of lyophilized sample. The protein quantification was carried out by Biuret method (NIELSEN, 2017)

### 2.3 Protein hydrolysates

To obtain protein hydrolysates, physiological human gastric fluids were simulated according to the methodology of MINEKUS *et al.* (2014). Prior to the ejecution, Simulated Gastric Fluid (SGF) and Simulated Intestinal Fluid (SIF) were prepared. For SFG, 25 mg of pepsin (2000 U/mg; MP Biomedicals, California, USA) was diluted in 50 mL of sodium chloride 0.35 M pH 2. For SIG, 61.6 mg of sodium hydroxide (Merck Millipore, Darmstadt, Germany) and 680 mg of sodium phosphate monobasic monohydrate (Sigma Aldrich, Missouri, USA) were dissolved in 100 mL of deionized water and adjusted to pH 7. For the gastric digestion, E:S mixture was used: 100 mg of the protein isolate was dissolved in 10 mL of NaCl 0.35 M pH 2.0. A relation 50:50 of diluted protein and simulated gastric fluid (SGF) was made for each sample. Subsequently, the samples were incubated during 2 hours at 37 °C and 500 rpm in a water bath (StableTemp, Cole Palmer, Illinois, USA). Immediately, a duodenal digestion was done, using pancreatin (Merck Millipore, Darmstadt, Germany) at a final concentration of 109 U/mL. A relation 1:1 (v/v) of the prior mixture and the simulated intestinal fluid (SIF) was made for each sample. Finally, the samples were incubated for 2 hours at 37 °C at 500 rpm in a water bath. Afterwards, the enzymatic reaction was stopped using 200 µL of sodium bicarbonate 1 M (Merck Millipore, Darmstadt, Germany) at 80 °C for 10 minutes. The samples were frozen at −80 °C and lyophilized. This protocol was carried out three times.

### 2.4 Antioxidant activities (TBARS assay)

TBARS assay (thiobarbituric acid reactive substances) is one of the most ancient and widely used assays in order to quantify oxidative stress (DASGUPTA *and* KLEIN 2014), which is based in the reaction between malondialdehyde and 2-thiobarbituric acid (TBA) that produces an adduct that shows pink color species, who absorbs at 532-535 nm. For this study, with minor modifications, the method of ROJANO, B. A., GAVIRIA, C. A. *and* SÁEZ (2008) was used. Aliquots of 0.2, 0.4, 1.0 and 2.0 mg of protein isolates or protein hydrolisates were mixed with 2 mL of distilled water (protein solution). Then, 500 µL of olive oil (previously oxidized) was mixed with 500 µL of protein solution. The samples were mixed at 450 rpm in a micro incubator Provocell Shaking (Esco, Singapore) for 13 hours, at 28 °C. Then, 1 mL TBA (1%) was placed and immediately incubated for 1 hour, at 95 °C. Finally, the samples were chilled at 4 °C for 15 minutes in a freezer and then analyzed at 532 nm in a UV Vis spectrophotometer (Thermo Scientific, USA). Butylated hydroxytoluene (BHT) ≥ 99% (Merck, Darmstadt, Germany), as positive control, was diluted in ethanol 96% (Merck, Darmstadt, Germany).

Oxidized olive oil was prepared with a heat treatment for 15 days. During the first eight days, the oil was heated in a laboratory oven (VWR, Pennsylvania, USA) at 70 °C. During latter seven days, the oil was heated at 40 °C. The results were compared with the positive control, quantifying the antioxidant activity percentage with this formula: % AA = (M – C) / C * 100; Where AA: Antioxidant activity, C: Oxidized oil absorbance (532 nm), M: Sample (protein sample + oxidized oil) absorbance (532 nm).

### 2.5 Anti-inflammatory activity

The anti-inflammatory activity of the protein isolates and hydrolisates was analyzed by following the methodology of PADMANABHAN P (2012), with minor modifications. Sodium diclofenac (25 mg/mL) was used as the positive control. Samples were homogenized, and aliquots of 0.2, 0.4, 1.0 and 2.0 mg were dissolved with 2 mL of deionized water. Then, the suspension was homogenized with 0.2 mL of egg albumin and 2.8 mL of phosphate buffered saline (PBS) pH 6.4. This suspension was mixed gently. The mixtures were incubated at 37°C for 15 minutes, then incubated at 70 °C for 10 minutes. The samples were finally chilled in cold water for 10 minutes, and the absorbance was analyzed using a UV-Vis spectrophotometer at 660 nm, with distilled water as a blank. The obtained data was compared with the results of the positive control (sodium diclofenac). The anti-inflammatory activity percentage was determined with this formula: % anti-inflammatory activity = (M-C)/C * 100, where C: Denatured egg albumin absorbance (without sample) and M: Sample absorbance (protein suspension + egg albumin).

### 2.6 Antimicrobial activity

In order to evaluate the antimicrobial activity, these certified strains were used: *Staphylococcus aureus* (ATCC^®^ 25923), *Escherichia coli* (ATCC^®^ 25922), *Bacillus cereus* (ATCC^®^ 10876), *Listeria monocytogenes* (ATCC^®^ 19115) and *Pseudomonas aeruginosa* (ATCC^®^ 10145). The guidelines of the Clinical and Laboratory Standards Institute (CLSI) were applied, with minor modifications. A culture of each bacteria was inoculated in Tryptic Soy Broth TSB (Merck Millipore, Darmstadt, Germany) until each culture’s optical density was 0.5 Mc Farland units. Then, in Mueller-Hinton agar plates (Oxoid, Basingstoke, Hampshire, UK), the inoculum was struck with a long cotton swab. The protein isolates and hydrolysates were evaluated at five concentrations: 500, 375, 250, 200 and 150 mg/mL, all diluted in distilled water. After the inoculum was struck in the agar, 6mm wells were made with sterile plastic pipette tips (200 µL). 70 µL of protein solution or controls were deposited into the 6mm wells. Gentamicin (500 µg/mL) was used as a positive control and sterile deionized water was used as a negative control. After the plates stood for 30 minutes, they were incubated for 48 hours, at 37 °C. The inhibition zones were measured after 24 hours and 48 hours. In order to determine the percentage of antimicrobial inhibition, this formula was used: % Inhibition = (B – A)/(C – A)*100. Where A: Negative control diameter (mm), B: Sample inhibition zone (mm), C: Positive control diameter (mm).

### 2.7 Statistical analysis

Statistical analysis was done by using Statgraphics Centurion 16.103 and Microsoft Excel^®^ software packages. Simple variance analysis (ANOVA) and Tukey’s comparative test at 95 % of confidence were used to evaluate statistical differences between protein extraction conditions and biological activities. Four (antioxidant activity) and three (anti-inflammatory and antimicrobial activities) replicates of each experiment were done, and the results were summarized as a mean and standard deviation.

## 3. RESULTS AND DISCUSSIONS

### 3.1 Preparation of protein isolates and hydrolizates

A proximate analysis of the baby lima bean flour was done at the National Institute of Agriculture Research of Ecuador (INIAP Santa Catalina). The analyzed samples had a mean of 21.15% of protein. This result is similar to the reported by FAO, 2005 (23.6%), TYLER, YOUNGS, & SOSULSKI, 1981 (22.8%) and SATHE, 2002 (17.5-28.7%).

Baby lima bean flour protein precipitation was based on isoelectric precipitation (CÁRDENAS *et al.* 2018). The pH was adjusted at four levels (3.0; 4.0, 5.0; 6.0) using HCl solution (2 N) and the precipitates were recovered. The highest percentage of protein was obtained at pH 5.0 (19.56 ± 1,55 %). At pH 6, no precipitation was observed, however the solution was again frozen and lyophilized. The precipitation effect was only present at pH <6. These precipitation effects could be related to the fact that a majority of legumes proteins have their isoelectric points under pH 6 (EL-SAYED M *et al.* 1986; KLUPŠAITĖ *and* JUODEIKIENĖ 2015).

The protein quantification of the isolates was determined by using the Biuret colorimetric method (DOREY *and* DRAVES 1998). At pH 5, protein isolate represents 62.53 ± 0,02 % (dry weight base). The protein isolates were characterized by Native-PAGE and SDS-PAGE (data pending for publication) which showed a predominance of 2S albumins and 11S globulins (Basic and acid subunities). Both gastric and intestinal digestibility were performed while simulating human physiologic conditions. The SDS-PAGE shows five protein bands with sizes between 15 kDa and 37 kDa. These results suggest that the proteins from baby lime beans were not completely hydrolyzed during human digestion. This effect was observed in related research, during which proteins from different legumes had a high persistence during proteases digestion (M CARBONARO, GRANT, *and* CAPPELLONI 2005; DESHPANDE & DAMODARAN 1989).

### 3.2 Antioxidant activity (TBARS method)

The antioxidant activity of protein isolates and hydrolysates were evaluated by the TBARS method. The protein isolates obtained at pH 3, 4, 5 and 6; and concentrations of 100, 200, 500 and 1000 µg/mL were evaluated (Table 1). The differences between precipitation pHs, concentrations and antioxidant activity were not statistically different (p < 0.05; simple variance analysis). However, at pH 5, the highest inhibition of lipid peroxidation was observed in all four of the evaluated concentrations (100, 200, 500, 1000 µg/mL), in relation to the obtained isolates at the pH 3, 4 and 6 levels.

**Table 1.** *In vitro* antioxidant activity of baby lima bean protein isolate using TBARS method.

| Precipitation pH | Concentrations |  |  |  |
| --- | --- | --- | --- | --- |
|  | 100 µg/ml | 200 µg/ml | 500 µg/ml | 1000 µg/ml |
| pH 3 | 10,82 ± 0,006<sup>a</sup> | 15,69 ± 0,007<sup>a</sup> | 22,30 ± 0,017<sup>a</sup> | 29,99 ± 0,012<sup>a</sup> |
| pH 4 | 13,97 ± 0,012<sup>a</sup> | 19,12 ± 0,011<sup>a</sup> | 23,81 ± 0,022<sup>a</sup> | 32,69 ± 0,008<sup>a</sup> |
| pH 5 | 15,66 ± 0,009<sup>a</sup> | 19,65 ± 0,008<sup>a</sup> | 24,45 ± 0,016<sup>a</sup> | 33,19 ± 0,024<sup>a</sup> |
| pH 6 | 13,40 ± 0,006<sup>a</sup> | 17,31 ± 0,009<sup>a</sup> | 22,60 ± 0,010<sup>a</sup> | 30,59 ± 0,027<sup>a</sup> |

Data are expressed as the mean ± Standard deviation SD (n=4).
Values in the same column having different letters differ significantly ( $p < 0.05$ ). ANOVA and Tukey's test

For this parameter, Butylated hydroxytoluene (BHT) ≥99% was used as a positive control at concentrations of 100, 200, 500 and 1000 µg/mL. Between 70,82 ± 0,042 (100 µg/mL) and 91,73 ± 0,050 % (1000 µg/mL), BHT showed considerable values of antioxidant activity. Figure 1 shows the relationship between protein isolates and hydrolysates at different pHs and the percentages of lipid peroxidation inhibition. The maximum antioxidant activity in protein isolates was observed at pH 5 and 1000 µg/mL, with a value of 33,19 ± 0,024 %. The protein hydrolysate obtained at pH 5 and 1000 µg/mL was at a value of 77,17 ± 0,029 % of lipid peroxidation inhibition. As expected, in comparison with the evaluated samples, the BHT showed considerable differences in their antioxidant activity.

**Figure 1.**
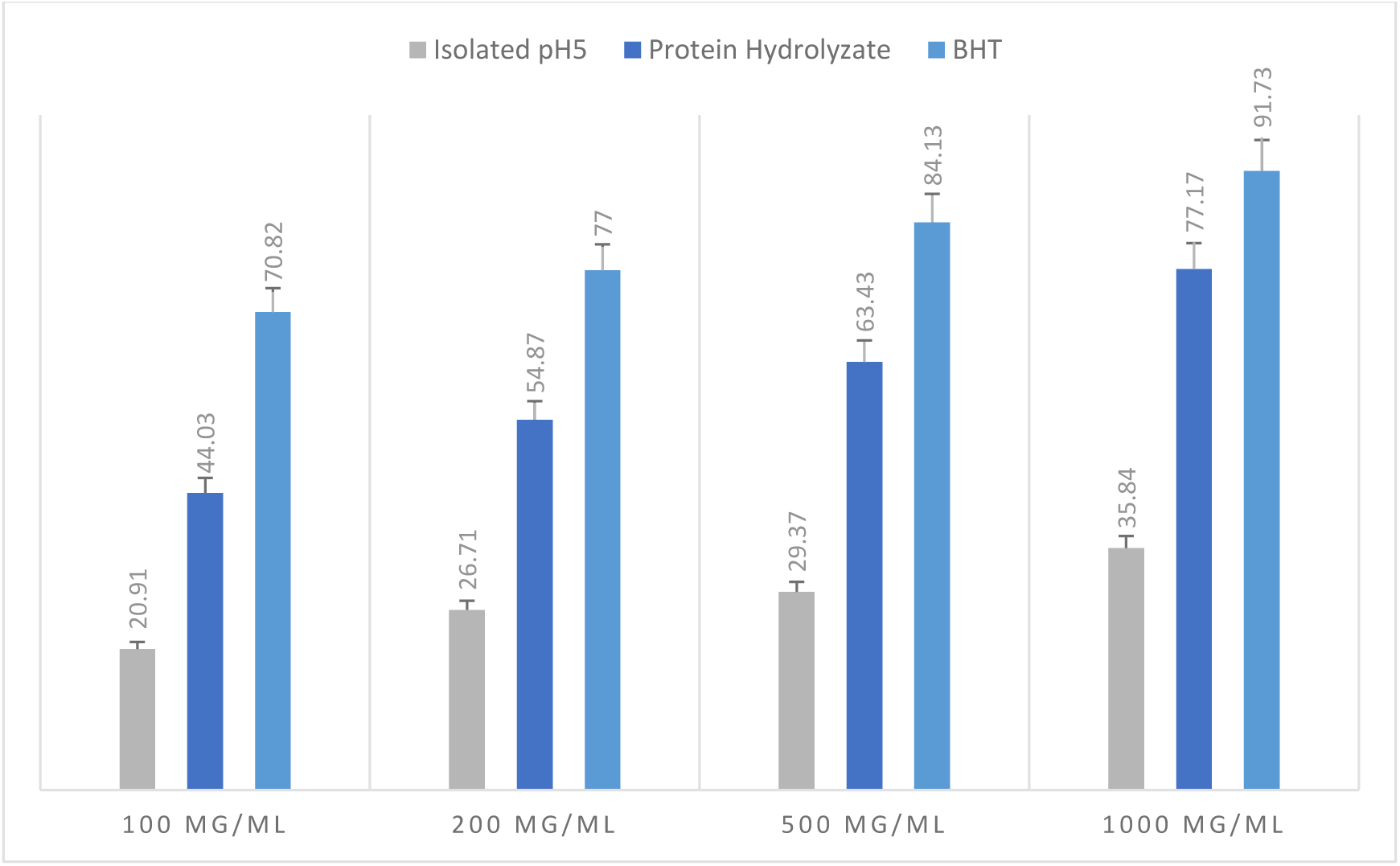
Inhibition of lipid peroxidation (TBARS) of the protein isolate and hydrolysate from baby lima bean at concentrations of 100, 200, 500 and 1000 μg/mL.

Figure 1 shows clear differences in antioxidant activity values in all of the evaluated concentrations as well as between isolates and hydrolysates while displaying statistical differences (p<0.05; simple variance analysis) between concentrations. In comparison with 100, 200 and 500 µg/mL, 1000 µg/mL was the highest observed peroxidation inhibition. This results supports studies which show an inverse correlation between molecular weight and antioxidant activity (ZHAO *et al.* 2012). The effect observed in different legume protein hydrolysates showed considerable antioxidant activity (DO EVANGELHO *et al.* 2016; DURAK *et al.* 2013; ALASHI *et al.* 2014; CARRASCO-CASTILLA *et al.* 2012).

### 3.3 *In-vitro* anti-inflammatory activities

Table 2 shows the anti-inflammatory activities of protein isolates obtained at four pH levels (3.0, 4.0, 5.0 and 6.0) and hydrolysates (pH 5). Diclofenac sodium (25 mg/mL) was used as a positive control at 100-1000 µg/mL. Statistical analysis do not show differences between anti-inflammatory activities and the precipitation pH of the isolates (p<0.05; simple variance analysis). However, for the protein isolates, the maximum activity was observed at pH 5 and 1000 µg/mL, with a value of 21,53 ± 0,39 % when inhibited by denatured protein.

**Table 2.** *In vitro* anti-inflammatory activity of baby lima bean protein isolate using albumin protein denaturation method.

| Precipitation pH | Concentrations |  |  |  |
| --- | --- | --- | --- | --- |
| | 100 $\mu\text{g/mL}$ | 200 $\mu\text{g/mL}$ | 500 $\mu\text{g/mL}$ | 1000 $\mu\text{g/mL}$ |
| pH 3 | $2,54 \pm 0,018^a$ | $4,00 \pm 0,038^a$ | $8,14 \pm 0,046^a$ | $10,14 \pm 0,038^a$ |
| pH 4 | $4,24 \pm 0,030^a$ | $7,36 \pm 0,032^a$ | $12,55 \pm 0,039^a$ | $17,52 \pm 0,056^a$ |
| pH 5 | $5,82 \pm 0,048^a$ | $11,09 \pm 0,054^a$ | $16,29 \pm 0,037^a$ | $21,53 \pm 0,039^a$ |
| pH 6 | $2,34 \pm 0,039^a$ | $5,42 \pm 0,033^a$ | $10,72 \pm 0,040^a$ | $15,80 \pm 0,057^a$ |

Data are expressed as the mean ± Standard deviation SD ( $n=4$ ).
Values in the same column having different letters differ significantly ( $p < 0.05$ ). ANOVA and Tukey's test

Protein hydrolysates (pH 5) showed an improvement in the percentages of inhibition in relation with protein isolates, with 30.62 ± 0.01 % of inhibition of denatured protein at 1000 µg/mL (Figure 2). Positive controls of diclofenac (100 – 1000 µg/mL) showed anti-inflammatory activities in the range of 32.14%–100.00%, as reported in related research (CÁRDENAS *et al.* 2018). Related research showed a clear inhibition of pro-inflammatory mediators of protein hydrolysates and unhydrolysed proteins from legumes (NDIAYE *et al.* 2012; POWNALL, UDENIGWE, *and* ALUKO 2010). Therefore, because baby lima bean proteins and hydrolysates have displayed an interesting anti-inflammatory activity, further research could be considered.

**Figure 2.**
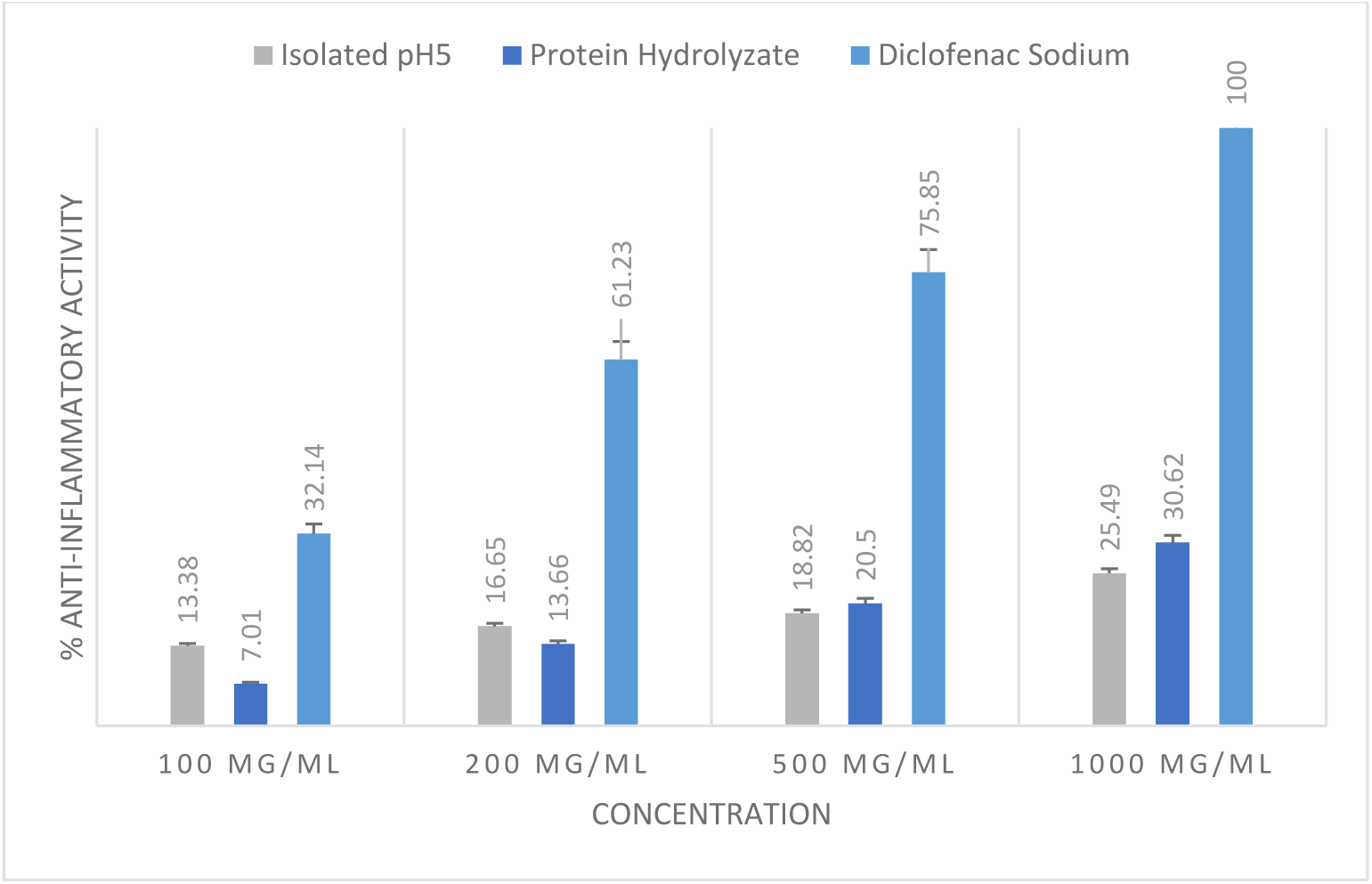
Anti-inflammatory Activity of protein isolate and hydrolysate from baby lima bean at concentrations of 100; 200; 500 and 1000 μg/ml.

### 3.4 Antimicrobial activity

Several protein hydrolysates from plant and animal origin, and their peptides, have shown antimicrobial activity against Gram-positive and Gram-negative bacteria. Other findings suggest that certain peptides can alter the cytoplasmic membrane or inhibit nucleic acid synthesis, protein synthesis or enzymatic reactions (BROGDEN 2005). However, the details of the peptides active mechanics have not yet been completely determined due to the vast number of peptides with potential biological activity (DASHPER, LIU, *and* REYNOLDS 2007).

To evaluate the potential antimicrobial activity of *P. lunatus* baby lima bean protein isolate and hydrolysates, the samples were screened against clinical bacterial strains (Figure 3). In the evaluated bacteria, between 150-500 mg/mL, no inhibition zones were observed in non-hydrolyzed protein samples. However, in four of five pathogenic bacteria evaluated (Table 3), their hydrolysates have shown a considerable antimicrobial activity. Significant differences were found among hydrolysate concentrations (p< 0.05; simple variance analysis).

**Table 3.** Bacterial inhibition areas from protein isolates and hydrolysates of *P. lunatus* baby lima beans against certified bacterial strains.

| Bacteria | Concentration | Isolate | Hydrolysate |  | Gentamycin<br>500 µg/ml (mm) |
| --- | --- | --- | --- | --- | --- |
|  |  | Inhibition<br>zone (mm) | Inhibition zone<br>(mm) | Inhibition (%) |  |
| *Escherichia coli*<br>(ATCC® 25922™) | 500 mg/ml | < 6 | 18,67 <sup>a</sup> ± 0,58 | 71,70 | 23,67 ± 0,58 |
|  | 375 mg/ml | < 6 | 15,67 <sup>b</sup> ± 0,58 | 54,72 |  |
|  | 250 mg/ml | < 6 | 12,33 <sup>c</sup> ± 0,58 | 35,82 |  |
|  | 200 mg/ml | < 6 | < 6 | 0 |  |
| *Pseudomonas aeruginosa*<br>(ATCC® 10145) | 500 mg/ml | < 6 | 12,33 <sup>a</sup> ± 0,58 | 33,31 | 25,00 ± 1,00 |
|  | 375 mg/ml | < 6 | 10,33 <sup>b</sup> ± 0,58 | 22,78 |  |
|  | 250 mg/ml | < 6 | 8,33 <sup>c</sup> ± 0,58 | 12,26 |  |
|  | 200 mg/ml | < 6 | < 6 | 0 |  |
| *Listeria monocytogenes*<br>(ATCC® 19115™) | 500 mg/ml | < 6 | 18,33 <sup>a</sup> ± 0,58 | 51,37 | 30,00 ± 1,00 |
|  | 375 mg/ml | < 6 | 16,33 <sup>b</sup> ± 0,58 | 43,04 |  |
|  | 250 mg/ml | < 6 | 13,67 <sup>c</sup> ± 0,58 | 31,96 |  |
|  | 200 mg/ml | < 6 | < 6 | 0 |  |
| *Bacillus cereus*<br>(ATCC® 10876) | 500 mg/ml | < 6 | 9,33 <sup>a</sup> ± 0,58 | 15,37 | 27,67 ± 0,58 |
|  | 375 mg/ml | < 6 | 7,33 <sup>b</sup> ± 0,58 | 6,14 |  |
|  | 250 mg/ml | < 6 | < 6 | 0 |  |
|  | 200 mg/ml | < 6 | < 6 | 0 |  |
| *Staphylococcus. aureus*<br>(ATCC® 25923™) | 500 mg/ml | < 6 | < 6 | 0 | 35,00 ± 1,00 |
|  | 375 mg/ml | < 6 | < 6 | 0 |  |
|  | 250 mg/ml | < 6 | < 6 | 0 |  |
|  | 200 mg/ml | < 6 | < 6 | 0 |  |

**Figure 3.**
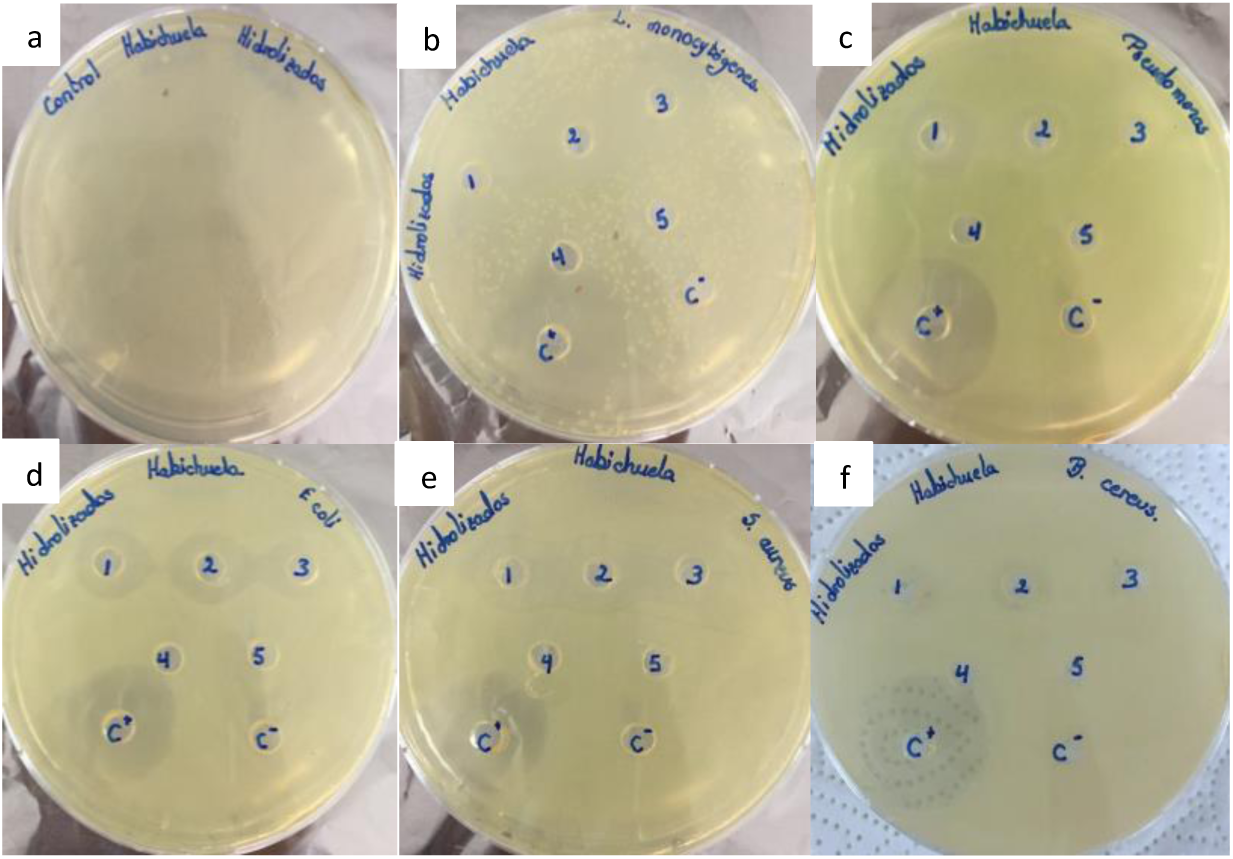
Antimicrobial activity of the protein hydrolisate obtained at pH 5. A) Protein hydrolysate without bacteria (negative control); B) L. monocytogenes; C) P. aeruginosa; in the D) E. coli; E) S. aureus; F) B. cereus.

The major percentage of inhibition was observed in 500 mg/mL of protein hydrolysate in Gram-negative *E. coli* ATCC ^®^ 25922, 71.7 % of which was related to the positive control (Gentamicin 500 µg/mL), followed by Gram-positive *L. monocytogenes* ATCC^®^ 19115 with 51.37 %. Gram-negative *P. aeruginosa* ATCC^®^ 10145 shows a considerable inhibition of 33.31 %. A slight inhibition of 15.37% was observed in Gram-positive *B. cereus* ATCC ^®^ 10876. The results were negative with Gram-positive *S. aureus* ATCC ^®^ 25923^™^. These results suggest that protein hydrolysates from *P. lunatus* baby lima beans possess a broad-spectrum of antimicrobial activity against Gram-negative and Gram-positive bacteria.

## 4. CONCLUSIONS

The *P. lunatus* baby lima bean protein isolates and hydrolysates displayed a considerable *in vitro* biological effect, based on the observed antioxidant, anti-inflammatory and antimicrobial activities. In comparison with protein isolates, protein hydrolysates performed better within the evaluated parameters. However, more in depth research is needed in order to evaluate the effect of bioactive peptides in baby lima bean proteins. Such peptides could be the source for notable biological activity for nutraceutical applications.

## ACKNOWLEDGMENTS

This work was supported by Dirección de Investigación y Desarrollo DIDE-Universidad Técnica de Ambato (Project 2074-CU-P-2016 “Valorización de la Calidad Nutricional de Alimentos Tradicionales de la Población Ecuatoriana VANFOOD”). This work have been reviewed in the English edition by Max Long BA AAS CELTA (UNSIS EFL Instructor, tenured).

## References

Alashi AM, Blanchard CL, Mailer RJ, Agboola SO, Mawson AJ, He R, Girgih A, Aluko RE, 2014. Antioxidant properties of Australian canola meal protein hydrolysates. Food Chem. 146:500–6.

Barrio DA, Añón MC, 2010. Potential antitumor properties of a protein isolate obtained from the seeds of Amaranthus mantegazzianus. Eur. J. Nutr. 49:73–82.

Bellucci E, Bitocchi E, Rau D, Rodriguez M, Biagetti E, Giardini A, Attene G, Nanni L, Papa R, 2014. Genomics of Origin, Domestication and Evolution of Phaseolus vulgaris. In: Genomics of Plant Genetic Resources: Volume 1. Managing, Sequencing and Mining Genetic Resources. pp 483–507.

Bernal G, Graham PH, 2001. Diversity in the rhizobia associated with *Phaseolus vulgaris* L. in Ecuador, and comparisons with Mexican bean rhizobia. Can. J. Microbiol. 47:526–34.

Betancur-Ancona D, Martínez-Rosado R, Corona-Cruz A, Castellanos-Ruelas A, Jaramillo-Flores ME, Chel-Guerrero L, 2009. Functional properties of hydrolysates from Phaseolus lunatus seeds. Int. J. Food Sci. Technol. 44:128–37.

Bitocchi E, Nanni L, 2012. Mesoamerican origin of the common bean (Phaseolus vulgaris L.) is revealed by sequence data. Pnas 109:788–96.

Brogden KA, 2005. Antimicrobial peptides: Pore formers or metabolic inhibitors in bacteria? Nat. Rev. Microbiol. 3:238– 50.

Broughton WJ, Hernández G, Blair M, Beebe S, Gepts P, Vanderleyden J, 2003. Beans (Phaseolus spp.) – model food legumes. Plant Soil 252:55–128.

Campos-Vega R, Guevara-Gonzalez RG, Guevara-Olvera BL, Dave Oomah B, Loarca-Piña G, 2010. Bean (Phaseolus vulgaris L.) polysaccharides modulate gene expression in human colon cancer cells (HT-29). Food Res. Int. 43:1057–64.

Carbonaro M, Grant G, Cappelloni M, 2005. Heat-induced denaturation impairs digestibility of legume (Phaseolus vulgaris L andVicia faba L) 7S and 11S globulins in the small intestine of rat. J. Sci. Food Agric. 85:65–72.

Carbonaro M, Maselli P, Nucara A, 2015. Structural aspects of legume proteins and nutraceutical properties. Food Res. Int. 76:19–30.

Cárdenas M, Carpio C, Welbaum J, Vilcacundo E, Carrillo W, 2018. Chia protein concentrate (Salvia hispanica l.) anti-inflammatory and antioxidant activity. Asian J. Pharm. Clin. Res. 11:382–6.

Carrasco-Castilla J, Hernández-Álvarez AJ, Jiménez-Martínez C, Jacinto-Hernández C, Alaiz M, Girón-Calle J, Vioque J, Dávila-Ortiz G, 2012. Antioxidant and metal chelating activities of peptide fractions from phaseolin and bean protein hydrolysates. Food Chem. 135:1789–95.

Dasgupta A, Klein K, 2014. Methods for Measuring Oxidative Stress in the Laboratory. Antioxidants Food, Vitam. Suppl.:19–40.

Dashper SG, Liu SW, Reynolds EC, 2007. Antimicrobial peptides and their potential as oral therapeutic agents. Int. J. Pept. Res. Ther. 13:505–16.

Debouck DG, Toro O, Paredes OM, Johnson WC, Gepts P, 1993. Genetic Diversity and Ecological Distribution of Phaseolus vulgaris (Fabaceae) in Northwestern South America. Econ. Bot. 47:408–23.

Deshpande SS, Damodaran S, 1989. Structure-Digestibility Relationship of Legume 7S Proteins. J. Food Sci. 54:108–13.

Dorey, Draves, 1998. Spectrophotometric determination of total protein-biuret Method. Quant. Anal. Lab. A New Approach Funded By Natl. Sci. Found.:1–3.

Durak A, Baraniak B, Jakubczyk A, Swieca M, 2013. Biologically active peptides obtained by enzymatic hydrolysis of Adzuki bean seeds. Food Chem. 141:2177–87.

El-Sayed M A-A. A, Ahmed S, Ahmed R E-M, Mohamed M Y, 1986. Extractability and Functional Properties of Some Legume Proteins Isolated by Three Different Methods. Sci. Food Agric. Volume 37:pages 553–559.

do Evangelho JA, Berrios JJ, Pinto VZ, Antunes MD, Vanier NL, Zavareze ER, 2016. Antioxidant activity of black bean (Phaseolus vulgaris L.) protein hydrolysates. Food Sci. Technol. 36:23–7.

FAO, 1995. Anexo 3: contenidos de nutrientes en alimentos seleccionados. Organ. Las Nac. Unidas Para La Aliment. Y La Agric.

Klupšaite D, Juodeikiene G, 2015. Legume: composition, protein extraction and functional properties. A review. Cheminê Technol. 1:5–12.

Minekus M, Alminger M, Alvito P, Ballance S, Bohn T, Bourlieu C, Carrière F, Boutrou R, Corredig M, Dupont D, Dufour C, Egger L, Golding M, Karakaya S, Kirkhus B, Le Feunteun S, Lesmes U, Macierzanka A, Mackie A, Marze S, McClements DJ, Ménard O, Recio I, Santos CN, Singh RP, Vegarud GE, Wickham MSJ, Weitschies W, Brodkorb A, 2014. A standardised static *in vitro* digestion method suitable for food – an international consensus. Food Funct. 5:1113–24.

Ndiaye F, Vuong T, Duarte J, Aluko RE, Matar C, 2012. Anti-oxidant, anti-inflammatory and immunomodulating properties of an enzymatic protein hydrolysate from yellow field pea seeds. Eur. J. Nutr. 51:29–37.

Ngoh Y-Y, Gan C-Y, 2016. Enzyme-assisted extraction and identification of antioxidative and α-amylase inhibitory peptides from Pinto beans (Phaseolus vulgaris cv. Pinto). Food Chem. 190:331–7.

Nielsen SS, 2017. Food Analysis. Springer.

Padmanabhan P JS, 2012. Evaluation of in-vitro antiinflammatory activity of herbal preparation, a combination of four herbal plants. Int J App Basic Med Sci 2(1):109–16.

Pownall TL, Udenigwe CC, Aluko RE, 2010. Amino Acid Composition and Antioxidant Properties of Pea Seed (Pisum sativum L.) Enzymatic Protein Hydrolysate Fractions. J. Agric. Food Chem. 58:4712–8.

El Productor, 2015. Ecuador: La habichuela gana terreno. Available from: https://elproductor.com/noticias/ecuador-la-habichuela-gana-terreno/

Rojano, B. A., Gaviria, C. A. Sáez JA, 2008. Determinación de la actividad antioxidante en un modelo de peroxidación lipídica de mantequilla inhibida por el isoespintanol. Vitae, 15(2):212–8.

Sathe SK, 2002. Dry bean protein functionality. Crit. Rev. Biotechnol. 22:175–223.

Tyler RT, Youngs CG, Sosulski FW, 1981. Air classification of Legumes. I. Separation Efficiency, Yield, and Composition of the Starch and Protein Fractions. Cereal Chem. 58:144–8.

Zhao Q, Selomulya C, Xiong H, Chen XD, Ruan X, Wang S, Xie J, Peng H, Sun W, Zhou Q, 2012. Comparison of functional and structural properties of native and industrial process-modified proteins from long-grain indica rice. J. Cereal Sci. 56:568–75.

